# Characterizing changes in land cover and forest fragmentation in Dhorpatan Hunting Reserve of Nepal from multi-temporal Landsat observations (1993-2018)

**DOI:** 10.1101/846741

**Authors:** Sandeep Sharma, Manjit Bista, Li Mingshi

## Abstract

Recent centuries have experienced drastic changes in land cover around the world where Himalayan countries like Nepal have undergone changes in the past several decades because of increasing anthropogenic pressure, natural risks and climatic factors. Accordingly, forest fragmentation has also been increasing alarmingly, which is a matter of concern for natural resource management agencies and biodiversity conservation communities. In this study, we assessed land cover change and forest fragmentation trends in Dhorpatan Hunting Reserve of Nepal by implementing landscape fragmentation and recovery process models, and calculating landscape indices based on five-date land cover maps derived from Landsat satellite images from 1993 to 2018. Six land cover types including forest, grass land, barren land, agriculture & built-up, water bodies and snow & glaciers were determined after an intensive field survey. Diverse derived image features were fed to the Support Vector Machines classifier to create land cover maps, followed by a validation procedure using field samples and reference data. Land cover maps showed an increase in forest area from 37.32% (1993) to 39.26% (2018) and snow & glaciers from 1.72% (1993) to 2.15% (2018) while a decrease in grassland area from 38.78% (1993) to 36.41% (2018) and agriculture & built-up area from 2.39% (1993) to 1.80% (2018). Barren land and water body showed negligible changes. The spatial explicit process of forest fragmentation indicated that shrinkage was the most responsible factor of forest loss while expansion was dominant to increment for forest restoration. High dependency of people persists on the reserve for subsistence resources being a cause of forest fragmentation and posing threats to biodiversity. Focus should be made on strategies to decrease the anthropogenic pressure on the reserve. This requires approaches that provide sustainable alternative resources to the local people and innovations that will help them become less reliant on natural resources.

## Introduction

Land use land cover (LULC) change is considered as one of the most important variables of global ecological change since its impacts are profound and range from global carbon and hydrologic cycle to aerosols and local biodiversity [1]. Land use refers to human activity on a piece of land for different purposes such as industrial zones, residential zones, and land cover refers to its surface features such as forest, grassland, etc [2, 3]. Land use is one of the main factors through which humans influence the environment. It involves both the manner in which the biophysical attributes of the land are manipulated and the intent underlying that manipulation [4]. There’s a direct link between land cover and the actions of people in their environment, i.e., land use may lead to land cover change.

There have been drastic changes in LULC over recent centuries where Nepal has undergone constant change over the past few decades due to major changes caused by anthropogenic and natural factors and their impacts on the national and regional environment and climate [5]. In Nepal, the Middle Mountain and High Mountain regions are more sensitive to LULC and are more seriously affected by it even with small changes, than the low land area [6]. In addition, forest fragmentation has been increasing alarmingly throughout the world [7], especially in the Hindu Kush Himalaya (HKH) region that has tremendous biodiversity, making it a major contemporary conservation issue. Forest fragmentation, is a process in which adjoining forest regions are increasingly being subdivided into the smaller patches (Gibson et al. 1988) which has negative effects on the ecosystems [8–11] including, increase in forest fire vulnerability, tree mortality, changes in species composition, seed dispersion and predation and easier access to the interior forest, leading to increased hunting and resource extraction [12, 13]. Therefore, it is a matter of worldwide concern and urgency for natural resource management agencies and biodiversity conservation communities to protect the forest resources upon which the health of the entire planet depends [14].

Land cover in the HKH region has undergone rapid change due to social, economic and environmental factors affecting the ecosystems and the services they provide [15]. The rate of degradation of forest land and the increase of cropland have been escalating since the 1970s which has created many environmental problems [16]. One of the ecologically important areas enclosed within the HKH is the Dhorpatan Hunting Reserve (DHR), which is also confronting increased anthropogenic and natural disturbances [17]. These disturbances have been intensifying the change of land cover along with fragmentation [18, 19]. Thus, understanding the forest fragmentation dynamics and LULC change patterns in DHR is urgently needed to plan future conservation strategies.

To characterize the land use change and forest fragmentation, studies based on multi-temporal Landsat images and landscape indices have previously been carried out in some mountainous protected areas which have similar properties to DHR [20, 21]. These studies discovered changes in land use and identified fragmentation as a result of anthropogenic disturbances. More studies related to LULC in the Himalayan areas of Nepal have been conducted in different areas over different time periods, for example, [22–27] and some research in DHR focused on distribution of species like common leopard (*Panthera pardus*), red panda (*Ailurus fulgens*) and blue sheep (*Pseudois nayaur*) [28–30], but none of them addressed LULC change and forest fragmentation in DHR. A new landscape fragmentation process model based on Forman’s theory was proposed by Li and Yang (31) to clarify the spatial process. Based upon this model, Ren, Lv (32) adapted a landscape restoration model to advance Forman’s theory and complete the characterization of landscape succession processes. These landscape fragmentation and restoration process model were used for forest landscape in the current study to fulfill the current limitation. This study was carried out to assist park managers and conservation partners to formulate future conservation policies and management strategies. The temporal and spatial changes observed in this study will help them generate interventions for future conservation. The main objectives of our study were; (1) to analyze the spatial-temporal trends of land cover change from 1993 to 2018, (2) to understand forest fragmentation trends with local biodiversity dynamics and (3) to valuate forest fragmentation by using fragmentation spatial process model, and landscape indices and follow up the socio-economic drivers for these changes.

## Methods

### Study area

Dhorpatan Hunting Reserve (DHR) is the only hunting reserve among 20 protected areas of Nepal. It extends from 28°15′ N to 28°55′ N latitude and 82°25′ E to 83°35′ E longitude covering an area of 1325 km^2^ (Fig 1). It was established in 1983 and was officially declared in 1987. The primary management objectives of the reserve are to allow hunting and to preserve the representative high altitude ecosystem with seven hunting blocks. Geographically it falls under the HKH region having altitudinal variation from 2000 meters to 7246 meters. The monsoon starts in June and lasts till October with an annual rainfall of 144.91 mm in 2018 [33].

**Fig. 1.**
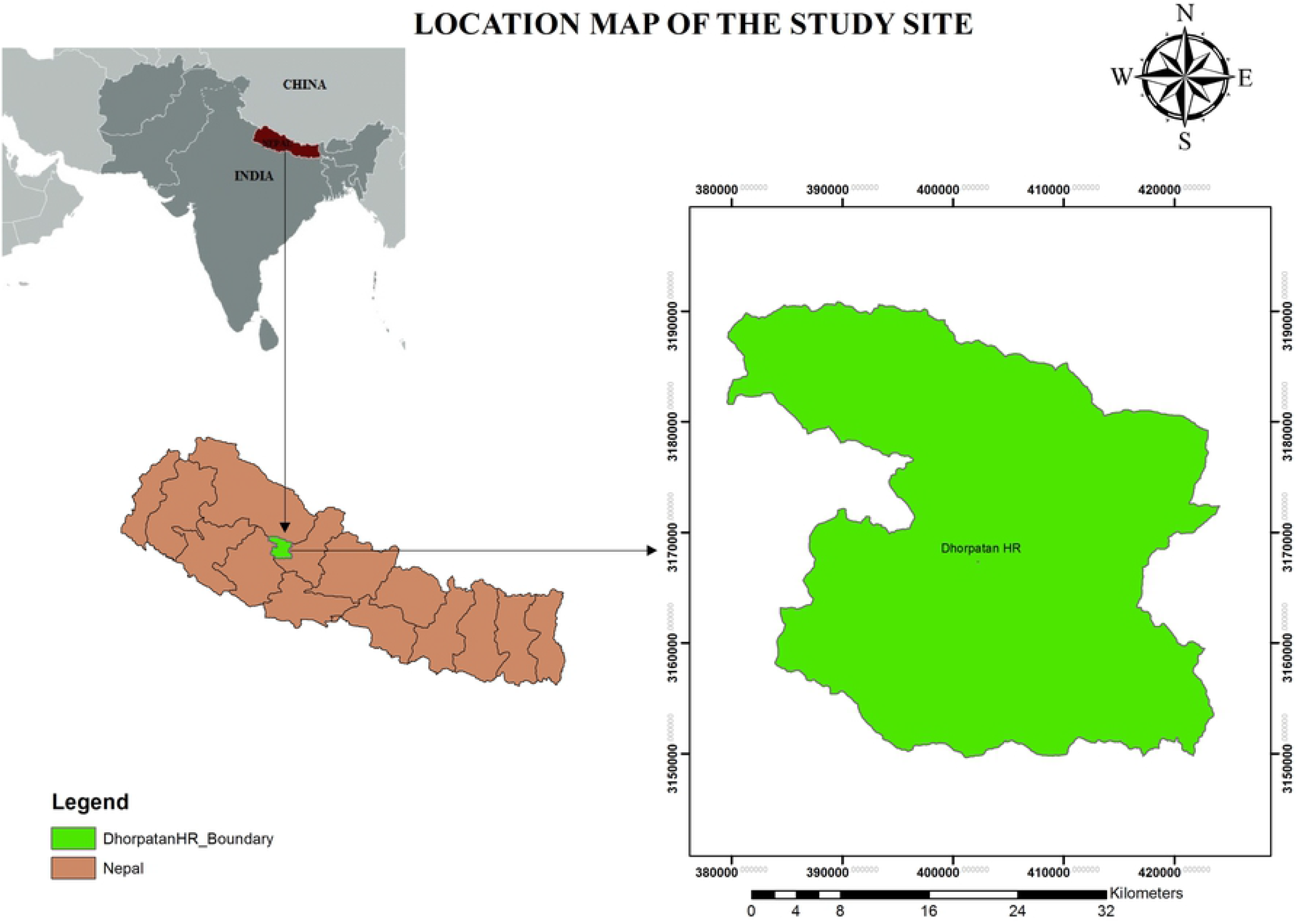
Location map of the study site.

DHR is comprised of high temperate, subalpine and alpine vegetation and has very high biodiversity values. The major tree species are *Pinus roxburghii, Taxus baccata, Pinus wallichiana*, and *Abies spectabilis* while the major faunal species are goral (*Nemorhaedus goral*), wild boar (*Sus scrofa*), Himalayan musk deer (*Moschus chrysogaster*), serow (*Capricornis sumatraensis*) and Indian muntjac (*Muntiacus muntjak*), leopard (*Panthera pardus*), lynx (*Felis lynx*), wild dog (*Cuon alpinus*), red fox (*Vulpes vulpes*), wolf (*Canis lupus*) and red panda (*Ailurus fulgens*) [34, 35]. The alpine flat pastures above treeline (4000m), locally known as *Patan* are very important for animals like blue sheep (*Pseudois nayaur*) which are preferred prey of *P. uncia*. The high elevation areas of the reserve mostly remain covered with cloud and higher altitude areas with snow. DHR is surrounded by human settlements except on the Northern side. There are around 47 small villages with 9,195 households and 43,078 population inside and around the reserve area [36].

## Data and Methods

### Data Source

Five cloud-free satellite images (Table 1) were downloaded from the United States Geological Survey (USGS) Center for Earth Resources Observation and Science (EROS) official website (http://earthexplorer.usgs.gov). Selection of the acquisition dates was driven by minimum cloud cover (or cloud contamination) to facilitate the monitoring of LC change in the study area. The downloaded remotely sensed data products were geometrically and atmospherically corrected to level 2 product type by USGS EROS data center, thus, their consistent and highest scientific standards and level of processing required for direct use in monitoring and accessing landscape change significantly reduced the magnitude of data processing for application users. Additionally, in view of the undulating mountainous terrain of the study area, we also downloaded DEM data with 30m resolution for the study area to support necessary terrain correction from the USGS website (https://earthexplorer.usgs.gov/). To support our computerized classification of LC, we paid an intense field visit to DHR during the period from September 28 to October 11, 2018 and collected 164 sample plots. We also collected 150 plots for 2011 and74 plots for 2004 from available previous national land use reference data.

**Table 1.** Details of Landsat data used in LC classification.

| Satellite | Sensor | Path/Row | Spatial<br>Resolution (M) | Acquisition<br>Date | Source |
| --- | --- | --- | --- | --- | --- |
| Landsat 5 | TM | 143040 | 30 | 09/11/1993 | USGS |
| Landsat 5 | TM | 143040 | 30 | 28/12/1999 | USGS |
| Landsat 5 | TM | 143040 | 30 | 22/10/2004 | USGS |
| Landsat 5 | TM | 143040 | 30 | 11/11/2011 | USGS |
| Landsat 8 | OLI | 143040 | 30 | 29/10/2018 | USGS |

### Image preprocessing

In view of the intense variability in elevation over the study area, prior to classifying the Landsat images, we implemented a SCS (Sun-Canopy-Sensor) + C-correction to minimize the negative terrain effects of rugged mountains on the actual pixels’ reflectance because this terrain correction strategy is rigorous, comprehensive and flexible and provides improved corrections compared to the SCS and four other photometric approaches (cosine, C, Minnaert, statistical-empirical) [37]. To further eliminate the effects of atmospheric interferences, derivation of NDVI, RVI and Greenness Index of Tasseled Cap Transform coupled with the post-terrain correction was done to support subsequent LC classifications.

### Image classification

The land cover classification scheme, including six major LC types (Forest, Grassland, Barren land, Water bodies, Agriculture (highly dispersed houses) and Snow & glaciers), was determined after a reconnaissance survey and discussion with local natural resource management experts. Built-up area was merged with Agriculture land because houses in the study area were scattered in small settlements and those settlements were further scattered which made it hardly possible to analyze separately by using the Landsat 30m resolution data only, without publicly available auxiliary geospatial data including multi-temporal auxiliary data sensitive to human activity patterns, such as distance to water, change in seasonal vegetation signals and distance to recent active fire [38]. Thus to minimize the complexity of classification, we merged the type of isolated houses into Agricultural type due to their few number of existence in the study region. Based on our field sample plots and other available auxiliary training data, using the derivatives mentioned in section 3.2, we implemented a support vector machines (SVM) classifier (the radial basis function was chosen as the kernel function, a cost parameter C to qualify the penalty of misclassification errors and γ were parameterized) to create time series land cover maps because SVM algorithm has the advantage of seeking an optimal solution to a classification problem and easy handling of a small number of samples over other machine learning algorithms such as decision tree and neutral networks [39].

### Accuracy assessment

Based on field validation samples and other auxiliary data (generally high spatial resolution Google Earth Maps and available historical dataset in Nepal), our created land cover maps were assessed by computing the overall accuracy, the kappa coefficient, the user accuracy, and the producer accuracy metrics from the confusion matrixes. However, due to the unavailability of the validation data in 1993 and 1999, we were unable to evaluate the land cover maps in these two years. We assumed that their classification reliability could be maintained because we applied same classifier to all the five-date images.

### Forest fragmentation and restoration process analysis

We followed Li and Yang (31) for the forest fragmentation spatial process analysis while for forest restoration process analysis, we followed Ren, Lv (32) to analyze its spatio-temporal characteristics. The entire analytical framework is summarized in Fig. 2. Specifically, through a spatial overlay operation of different time periods classified maps, we acquired information about the lost and remaining patches of interesting landscape elements. Then the lost patches were reclassified into four different categories based on their spatial relationship with the remaining patches. The lost patches surrounded by remaining patches were classified as perforation; those connected with two or more remaining patches were classified as subdivision. If the lost patches were connected with only one remaining patch then they were categorized under shrinkage and finally, those isolated from remaining patches were categorized as attrition. Forest restoration analysis based on Ren, Lv (32) (Fig 2.ii) model was applied based on two spatial processes: increment and expansion. Increment referred to those restorative patches which were isolated from the remaining forest patches and expansion meant the patches were connected to one or more remaining forest patches. After generating new/remaining forest pixel, we got the new/remaining forest layer. Focal variety statistics accompanying the Moore neighborhood method was applied to the new forest cell to identify different remnant patches in its eight neighboring cells. Then Zonal maximum statistics analysis was used to identify the maximum neighborhood variety value for all newly gained forest patches. The return values were 1 and 2 which were classified as increment and expansion respectively. The fully analytical models were developed and implemented in ArcToolbox model builder environment.

**Fig. 2.**
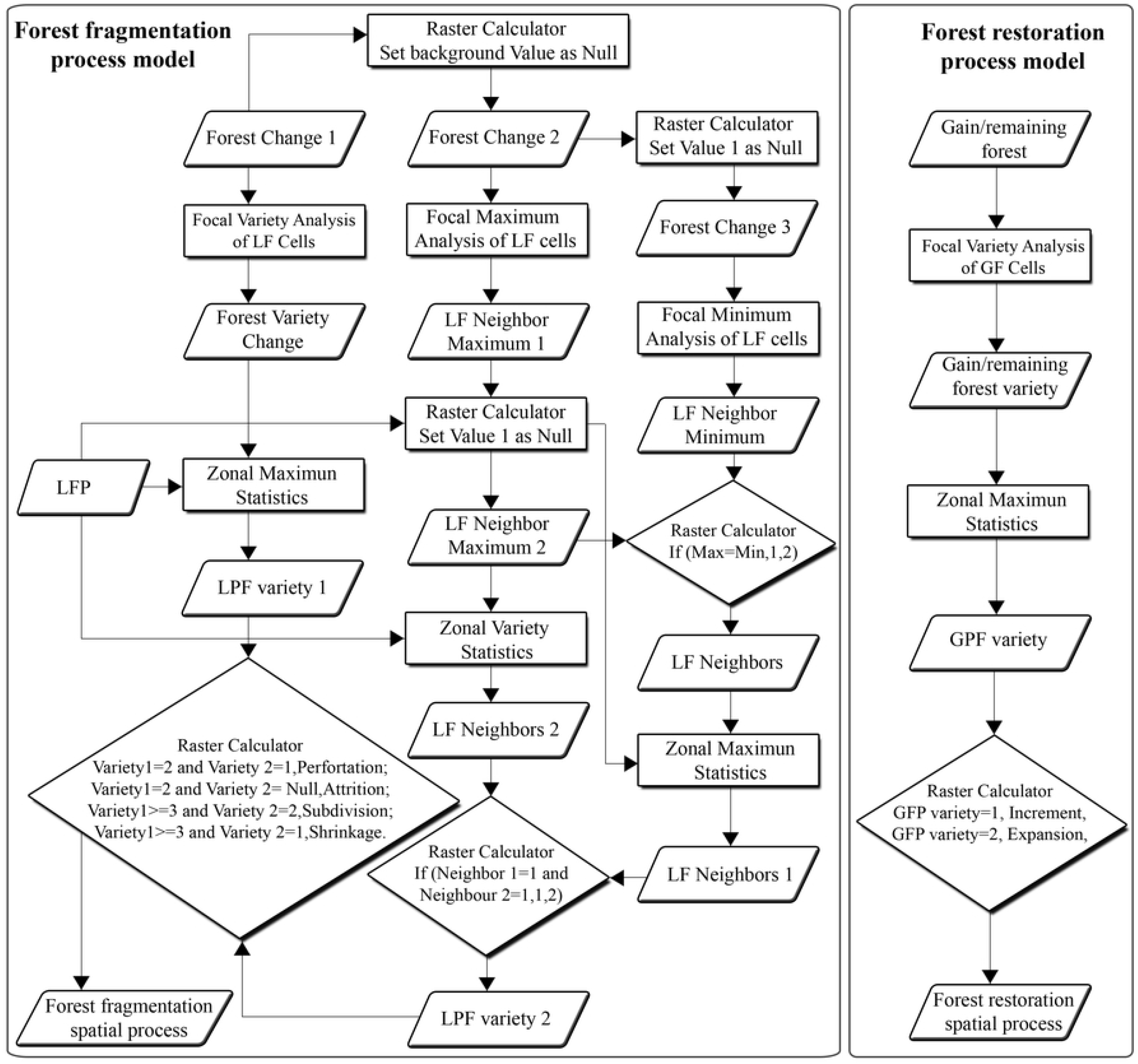
i) Conceptual model of four spatial processes of landscape fragmentation and ii) forest restoration model.

### Landscape metrics analysis at the class level

Unlike the fragmentation and restoration models, which produce spatially explicit maps showing forest changes over time and space, the landscape metrics computation can provide statistics describing landscape change trends quantitatively. FRAGSTATS 4.4 is a spatial pattern analysis program that quantifies the composition and configuration of the patches and provides information on the structure and spatial pattern of landscape [40], which calculates indices characterizing each patch in the mosaic, each patch class in the mosaic and the landscape mosaic as a whole. In the current work, we calculated largest patch index (LPI), edge density (ED), perimeter area fractal dimension (PAFRAC), and aggregation index (AI) at the class level, with an emphasis on forest class.

## Results

### Accuracy assessment

Table 2 displays the user’s accuracy, producer’s accuracy and kappa coefficients of three time periods. The overall accuracy for both the years, 2018 and 2011 was 92% but for 2004 was 84%. We did not have adequate quality data (high spatial resolution Google Earth Maps and available historical dataset) for the year 2004 to get a higher overall accuracy. The calculated Kappa coefficients were 88%, 87%, and 81% respectively for the years 2018, 2014 and 2004 which meant that the level of agreement was strong.

**Table 2.** Accuracy assessment result for 2018, 2011 and 2004

| Land Cover | Year |  |  |  |  |  |
| --- | --- | --- | --- | --- | --- | --- |
|  | 2018 |  | 2011 |  | 2004 |  |
|  | User's<br>accuracy | Producer<br>accuracy | User's<br>accuracy | Producer<br>accuracy | User's<br>accuracy | Producer<br>accuracy |
| Forest | 0.98 | 0.95 | 1.00 | 0.92 | 0.94 | 0.78 |
| Snow & glacier | 1.00 | 1.00 | 1.00 | 0.67 | 1.00 | 0.92 |
| Agriculture and<br>built-up area | 0.75 | 1.00 | 1.00 | 1.00 | 0.40 | 0.80 |
| Grass land | 0.81 | 0.95 | 0.82 | 0.97 | 0.88 | 0.88 |
| Water bodies | 1.00 | 1.00 | 1.00 | 0.67 | 0.50 | 0.50 |
| Barren land | 0.98 | 0.86 | 0.85 | 0.88 | 0.83 | 0.83 |
| Overall accuracy | **0.92** |  | **0.92** |  | **0.84** |  |
| Kappa Coefficient | **0.88** |  | **0.87** |  | **0.81** |  |

### Land cover change

The study area has been classified into six different land cover classes as shown in Fig. 3 for the years 1993, 1999, 2004, 2011 and 2018. It is apparent from the five maps that the major portion of DHR is covered by forest, grass land and barren land. Forest has occupied the Western and the Southern part of the study area while grass land has spread from the middle of the reserve towards the North, between forest and barren land. Snow & glacier have occurred in the Northeast part while agriculture and built-up are distributed in the lower altitude near the forest. The changes in the six land cover classes on different time point maps are not distinctly visible as the percentage of changes were significantly low.

**Fig. 3.**
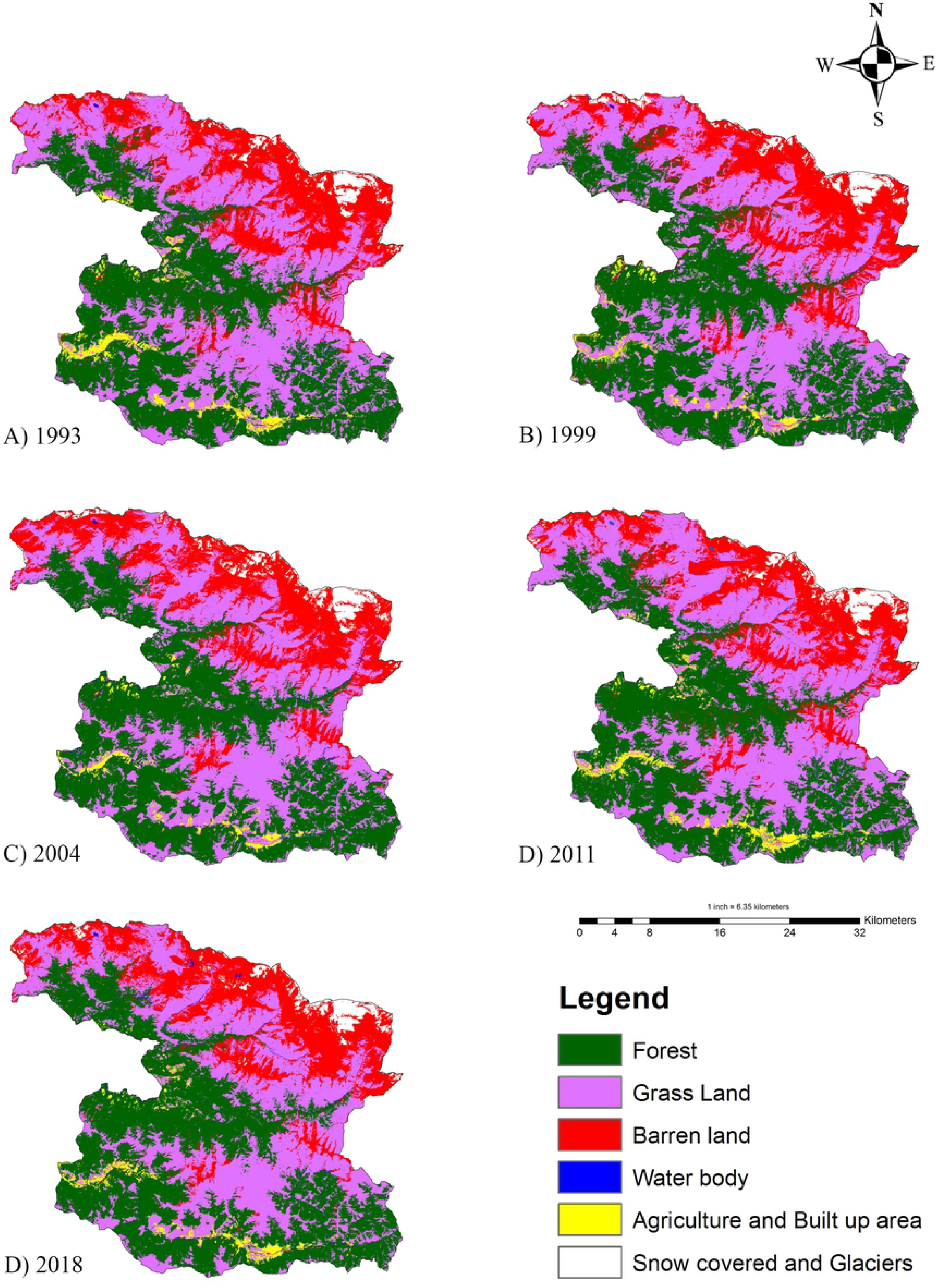
Supervised classified land cover maps of five different study time periods.

The analysis of Table 3 shows that the forest and grass land together covered around 76% of the landscape during all years. Barren land being the third largest LC class covered around 20% area during all time periods. Water, agriculture & built-up and snow & glaciers respectively covered around 0.5%, 2% and 2% on average during the five time points.

**Table 3.** Area and percentage of different land cover class in km^2^ from the year 1993–2018.

| Land cover distribution |  |  |  |  |  |  |  |  |  |  |
| --- | --- | --- | --- | --- | --- | --- | --- | --- | --- | --- |
| LC categories | 1993 |  | 1999 |  | 2004 |  | 2011 |  | 2018 |  |
|  | Area | Area | Area | Area | Area | Area | Area | Area | Area | Area |
|  | (km<sup>2</sup>) | (%) | (km<sup>2</sup>) | (%) | (km<sup>2</sup>) | (%) | (km<sup>2</sup>) | (%) | (km<sup>2</sup>) | (%) |
| **Forest** | 494.52 | 37.32 | 499.00 | 37.66 | 512.01 | 38.64 | 483.13 | 36.46 | 520.23 | 39.26 |
| **Grass land** | 513.90 | 38.78 | 517.22 | 39.04 | 482.62 | 36.42 | 524.12 | 39.56 | 482.37 | 36.41 |
| **Barren land** | 256.11 | 19.33 | 256.30 | 19.34 | 274.08 | 20.69 | 251.48 | 18.98 | 264.51 | 19.96 |
| **Water body** | 6.23 | 0.47 | 4.54 | 0.34 | 6.48 | 0.49 | 7.38 | 0.56 | 5.64 | 0.43 |
| **Agriculture & built-up** | 31.68 | 2.39 | 19.74 | 1.49 | 18.73 | 1.41 | 30.14 | 2.27 | 23.90 | 1.80 |
| **Snow & glacier** | 22.77 | 1.72 | 28.34 | 2.14 | 31.29 | 2.36 | 28.95 | 2.18 | 28.55 | 2.15 |
| **Total** | 1325.00 | 100.00 | 1325.00 | 100.00 | 1325.00 | 100.00 | 1325.00 | 100.00 | 1325.00 | 100.00 |

The LC change percentages in all five time periods depict that the changes were very low, the uttermost change not exceeding 3.5% in any time period for any LC class. The greatest change in the entire study period was observed in grass land which decreased by 2.38%. Conversely, forest cover had increased by 1.94% over the span of 25 years. Agriculture & built-up area showed a decrease of 0.59% while snow & glacier increased by 0.44%. Water showed a negligible decrease of 0.04% while barren land had an increase of 0.63% throughout the entire study period. It can be inferred that grass land has been converted to barren, snow & glacier and forest. Table 4 summarizes the specific transition information for the different time periods and the whole time interval studied.

**Table 4.**
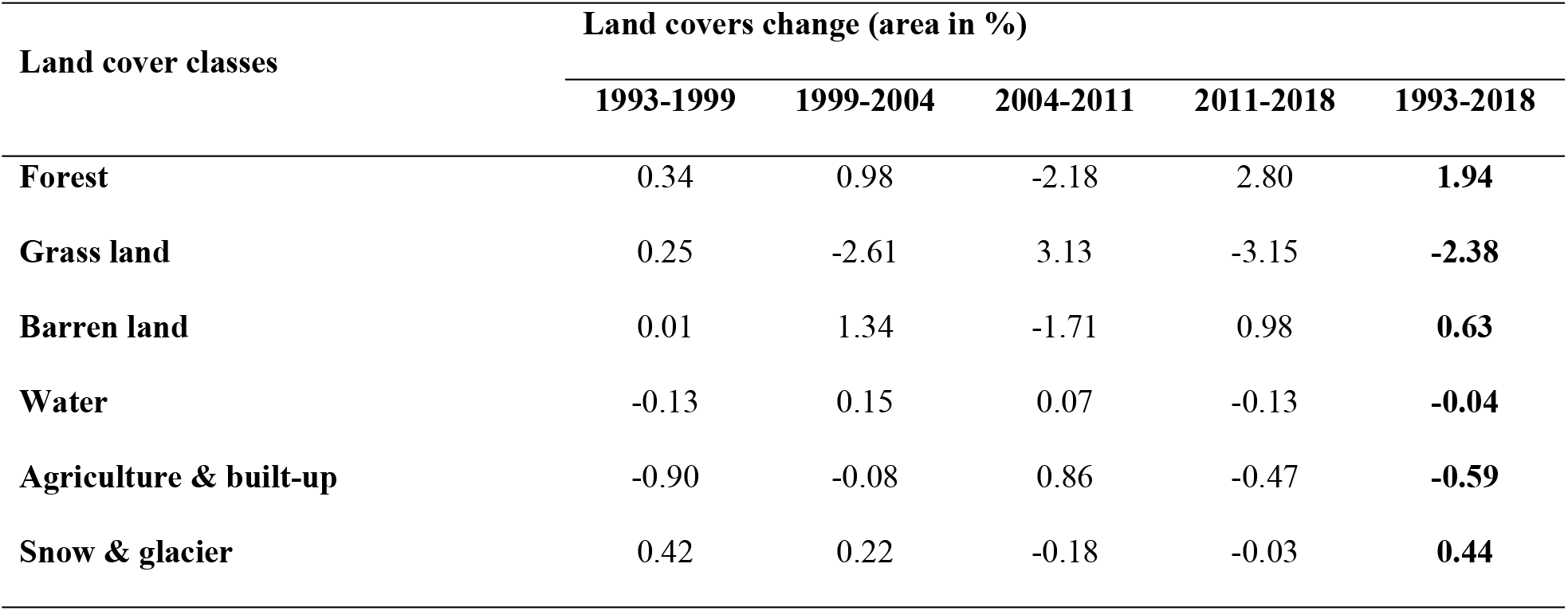
Change in percentage of different land cover classes during 1993–2018 (positive values indicate increase while negative values indicate decrease)

| **Land cover classes** | **Land covers change (area in %)** |  |  |  |  |
| --- | --- | --- | --- | --- | --- |
|  | **1993-1999** | **1999-2004** | **2004-2011** | **2011-2018** | **1993-2018** |
| **Forest** | 0.34 | 0.98 | -2.18 | 2.80 | **1.94** |
| **Grass land** | 0.25 | -2.61 | 3.13 | -3.15 | **-2.38** |
| **Barren land** | 0.01 | 1.34 | -1.71 | 0.98 | **0.63** |
| **Water** | -0.13 | 0.15 | 0.07 | -0.13 | **-0.04** |
| **Agriculture & built-up** | -0.90 | -0.08 | 0.86 | -0.47 | **-0.59** |
| **Snow & glacier** | 0.42 | 0.22 | -0.18 | -0.03 | **0.44** |

### Forest fragmentation and restoration spatial process analysis

The spatially explicit process of forest change captured by the forest fragmentation and restoration process models are depicted in the five time period maps below (Fig. 4). The maps show how the fragmented and restored patches are irregularly dispersed over DHR. Among the four different categories of fragmentation, perforation occurred mostly in the core forest patches while subdivision distributed near the edge of large forest patches. Shrinkage occurred from the core forest area to the medium-sized forest patches whereas attrition scattered over the small forest patches. Shrinkage on average was the most responsible and common spatial process for loss of forest patches in terms of area which was followed by subdivision. In case of restoration, both the increment and the expansion were dispersed around the whole study area but were more concentrated on the southern side. Expansion covered significantly larger area than the increment where the greatest expansion occurred during the period 1999-2004 (54.08 km^2^) which was closely followed by the period 2011-2018 (53.59 km2).

**Fig. 4.**
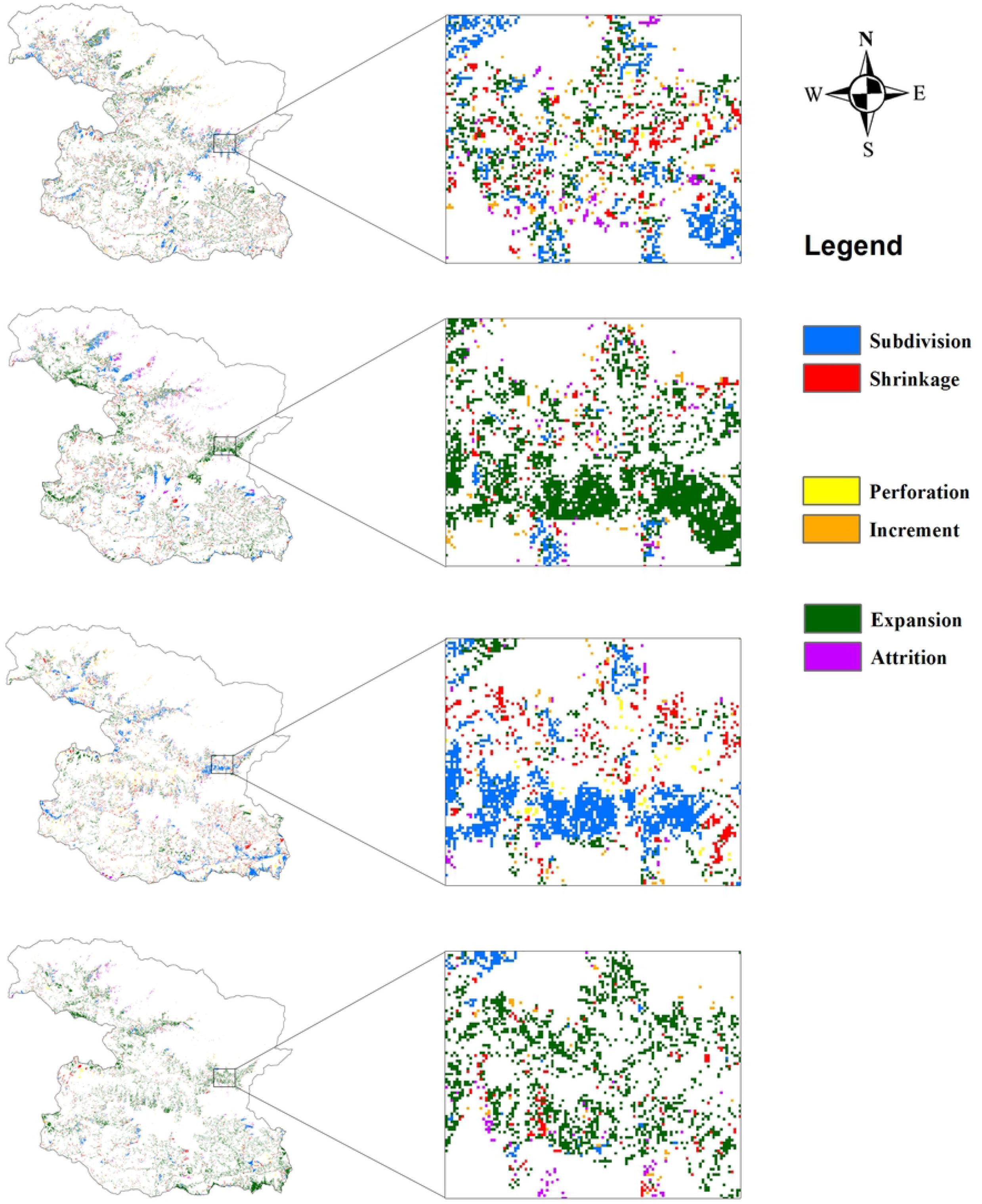
Forest fragmentation and restoration spatial process in DHR during the period 1993 to 2018.

Table 5 displays the area and percentage of forest land generated by four different fragmentation spatial processes and two restoration spatial processes. During the three periods; 1993-1999, 1999-2004 and 2011-2018, shrinkage contributed as the foremost spatial process for forest area loss which accounted for 49.04%, 44.28% and 50.74% respectively. For these same three time periods subdivision was the second major process for forest loss with 39.26%, 36.73% and 25.74% respectively. For the interval of 2004-2011, subdivision was the most responsible process for forest loss with 44.60% followed by shrinkage with 39.66%. The perforation and attrition explained little forest loss as per fragmentation spatial process analysis during all time periods. In terms of forest area loss, the period from 2004 to 2011 experienced a loss of nearly 54 km2 while the period from 2011-2018 only covered 24.28 km2. The restoration process model of different time periods revealed that more than 54 km^2^ area of forest patch expansion was gained in 1999-2004 and nearly the same area (53.59 km2) area of expansion occurred in 2011-2018. The expansion mostly occurred in the southeast and northwest parts of the hunting reserve and was dominant to increment over the whole study period.

**Table 5.** Forest fragmentation spatial explicit and restoration process analysis of four-time periods.

| Time period → | 1993-1999 |  | 1999-2004 |  | 2004-2011 |  | 2011-2018 |  |
| --- | --- | --- | --- | --- | --- | --- | --- | --- |
| Six Spatial process | Km<sup>2</sup> | % | Km<sup>2</sup> | % | Km<sup>2</sup> | % | Km<sup>2</sup> | % |
| **Fragmentation** |  |  |  |  |  |  |  |  |
| Perforation | 2.11 | 4.63 | 1.55 | 3.51 | 6.32 | 11.82 | 1.36 | 5.60 |
| Subdivision | 17.91 | 39.26 | 16.2 | 36.73 | 23.84 | 44.60 | 6.25 | 25.74 |
| Shrinkage | 22.37 | 49.04 | 19.53 | 44.28 | 21.2 | 39.66 | 12.32 | 50.74 |
| Attrition | 3.23 | 7.08 | 6.83 | 15.48 | 2.09 | 3.91 | 4.35 | 17.92 |
| Total | 45.62 | 27.24 | 44.11 | 26.34 | 53.45 | 31.92 | 24.28 | 14.50 |
| **Restoration** |  |  |  |  |  |  |  |  |
| Increment | 4.93 | 9.93 | 1.72 | 3.08 | 2.57 | 9.38 | 1.98 | 3.56 |
| Expansion | 44.7 | 90.07 | 54.08 | 96.92 | 24.82 | 90.62 | 53.59 | 96.44 |
| Total | 49.63 | 26.34 | 55.80 | 29.62 | 27.39 | 14.54 | 55.57 | 29.50 |

A close scrutiny of Table 5 shows that the degree of fragmentation was similar for the first two time periods but increased significantly in the 3^rd^ time period (2004-2011) and finally dropped down in the last time period (2011-2018). The restoration on the other hand increased slightly on the 2^nd^ time period (1999-2004), then plunged sharply for the 3^rd^ period (2004-2011) and retrieved on the 4^th^ period as equivalent to the 2^nd^ period. Overall, the area of forest gained from restoration was nearly 21 km^2^ more than the area lost from forest fragmentation.

### Landscape metrics analysis at class level

Table 6 presents the landscape metric analysis at the class level for forest cover class. The values of largest patch index (LPI) ascended to the highest 32.42% in 1999. Except in this year, the total landscape area comprised by the largest patch of forest was significantly low ranging from 16.16% to 18.64%. Edge density (ED), an important variable to measure forest fragmentation, was noted highest for the year 1993 (46.19 m/ha). The values were comparatively higher in the years 1999 (45.99 m/ha) and 2011 (44.55 m/ha) but was lower in 2018 (36.70 m/ha) and the lowest in 2004 (34.33 m/ha). This showed that less fragmentation occurred in 2004 and 2018 than in other three time points. The Perimeter-area fractal dimension (PAFRAC) increased in patch shape complexity, approaching 2, which means that the shape is highly complex. The average value for PAFRAC was 1.46 for all years, least value being 1.42 in 2004 and 2018 and highest value being 1.50 in 1999. The aggregation index (AI) value showed that around 91.72% of the forest cover class was adjacent to other different classes in average. AI increases as the focal patch type is increasingly aggregated and equals 100 when the patch type is maximally aggregated into a single, compact patch. In summary, results from all the four metrics show a similar trend of fragmentation where the years 2004 and 2018 experienced less forest fragmentation compared with other time points.

**Table 6.** Landscape metric analysis at the class level for forest cover class.

| Class Metrics (Unit) | Forest cover class |  |  |  |  |
| --- | --- | --- | --- | --- | --- |
|  | Year |  |  |  |  |
|  | 1993 | 1999 | 2004 | 2011 | 2018 |
| **LPI (%)** | 16.25 | 32.42 | 18.64 | 16.16 | 18.09 |
| **ED (m/ha)** | 46.19 | 45.99 | 34.33 | 44.55 | 36.70 |
| **PAFRAC (no unit)** | 1.49 | 1.50 | 1.42 | 1.47 | 1.42 |
| **AI (%)** | 90.68 | 90.82 | 93.31 | 90.83 | 92.98 |

## Discussion

### Land cover change

The land cover of DHR has been changing slowly from 1993 to 2018 as displayed in Table 4. The overall trend revealed that two larger sections (forest and grassland) have the highest change rates compared to other land cover classes. In this study area, forest cover has increased by 1.94% with an annual rate of 0.08% which is consistent with the national forest growth of Nepal; i.e. 5.14% from 1994 to 2014 [41]. Effective enforcement of laws, mobilization of the security forces and fulfillment of reserve staff curbed illicit collection of forest products and movement of unwanted visitors [42] which assisted to enrich the forest. It is possible that the treeline has shifted a little higher with the global warming as observed in the study in central Nepal where *Abies spectabilis* has shifted 2.61 m year^−1^ upward in Manaslu Conservation Area [43].

Grassland, on the other hand, has decreased by 2.38% with an annual rate of 0.10% throughout the study period. Other studies conducted in the highlands of Nepal have discovered similar results, for example [5, 44]. One important reason for grassland shrinkage may be trampling by the livestock movement in the reserve as explained by Panthi, Khanal (45) which not only reduces forage ability but also compacts the soil and leads to gully formation.

The results displayed in Table 4 clearly shows that the changes in the barren land, water, agricultural land and snow & glaciers are comparatively less during all the five time periods. An overall decrease is observed in the agricultural and built-up area while the opposite is the case for the snow & glaciers. Most of the agricultural lands situated in High Mountains have been abandoned by the people in recent years [46] due to labor shortages, a consequence of youth emigration. These lands have been slowly converting into shrub land which will eventually develop to forest. Overall, the agriculture land and built-up area have decreased but encroachment problem in this reserve exists in some accessible areas. According to a report from the reserve, 175-hectares of land have been encroached by people migrating from a more rural part of the reserve [42].

For the snow cover & glaciers of DHR, slight increase of 0.44% with an annual rate of 0.02% was observed from 1993 to 2018. Different researches on climate change have shown rapid change of snow cover and glacier meltdown. A study in Langtang valley of Nepal, whose geographic properties are alike to DHR, showed a decrease of 26% in snow-covered land from 1977 to 2010 [47] while another review study by Paudel, Zhang (5) over the whole country showed an increase of 4.61% from 1979 to 2010. One possible reason for the fluctuation could be the climatic or seasonal variation (temporal cover of snow) over the time period.

### Forest fragmentation and local biodiversity dynamics

Forest fragmentation occurs through the intensification of anthropogenic activities [12]. However in this study, although many human settlements existed in and around DHR and the local people highly depended on the forest for various resources, fragmentation in DHR was not so severe but was irregularly distributed on four out of seven hunting blocks of the reserve, namely, *Surtibang*, *Flagune and Barse blocks* in the southern part, and *Seng* block in the northern part. The result of forest fragmentation spatial process analysis showed that shrinkage was the most common fragmentation process followed by the subdivision. The result from the class metric analysis indicated that ED has increased from 1993 to 2011 and decreased in 2018. ED is very important in understanding the spatial heterogeneity as higher ED means higher spatial heterogeneity [40]. ED in DHR has high values ranging from around 34.33 m/ha to 46.19 m/ha, which means forest patches were distributed in small patches and/or having irregular shapes [48]. A similar case was observed in Dudhawa National Park of India where Midha and Mathur (49), reported that higher ED value is caused by fragmentation. In this study in DHR most of the forest patches were all connected with other lands.

Fragmentation is one of the most important factors which tend to have negative impact on continuity and quality of forests [50], loss of habitat [51], loss of biodiversity [52, 53], ecosystem functions [54] and facilitates the establishment of invasive species exposing the endemic plants and animals to vulnerability [55]. DHR is one of the prime habitats of the endangered fauna like the *A. fulgens*. Its habitat is characterized by the presence of mixed deciduous and coniferous forest [56], with more preference for forest with ringal bamboo (*Arundinaria spp*) understory. Trees species like Abies *spp*, oaks, and *Pinus spp* have been increasingly eliminated from the fragmented region to meet timber demand of local communities. This could be detrimental to *A. fulgens* since these tree species provide important resting and nesting cover for the small mammal [45]. Some mammal species need large core areas of forest as their primary habitat [57], and these species might be affected in DHR due to fragmentation.

These impacts related to forest fragmentation which are posing threat to biodiversity can be impaired by introduction of adaptive interventions for ecological restoration. This is achievable by enhanced understanding of the dynamics of land cover change and forest fragmentation [58, 59]. Park management and conservation partners can benefit from the result of this study to recuperate the observed patches. Recovery of fragmentation is necessary not only for biodiversity protection but also to control the possible conflict between humans and wildlife because forest fragmentation certainly becomes a critical driver of such conflicts [60].

### Socioeconomic drivers responsible for the observed changes

Studies have shown that socioeconomic factors have been responsible for forest fragmentation when people are extremely dependent on forest resources, like in Romania [61], in India [62] and in Bolivia [63]. DHR has substantial human presences that exceedingly rely on forest resources for their daily livelihood. Around 47 small villages are scattered inside the reserve and many others exist at the periphery. The number of households (5,568) and residents (35,310) in 2007 [17] has increased to (9,195) and (43,078) respectively in 2011 [36] with around 98% of the HHs using firewood as the main source of energy. Fodder, forage, timber, bamboo, medicinal herbs, etc, are the other major resources extracted from the reserve. Thapa, Panthi (64) discovered that around 4000-5000 people go to the alpine and subalpine pastures of DHR from late spring to early summer in search of the lucrative Yarsagumba (*Ophiocordyceps sinensis)*. Anthropogenic pressure is growing on the reserve (both in the pasture and forest) due to the availability of Yarsagumba.

Along with traditional agricultural, the major sources of livelihood of local people of this region are animal husbandry and trans-boundary trade [65]. Unplanned and unsystematic grazing widespread across the area has contributed in suppression of new regeneration, which eventually has aided in forest fragmentation. Some local herdsmen intentionally set fire each year to promote the growth of herbaceous vegetation for livestock forage which sometimes can convert to uncontrolled forest fire leading to greater catastrophe. The number of livestock, though decreased by around 1,14,104 in 2007 to 78,025 in 2011, the pastures of DHR is still used extensively for grazing.

This study showed that forest fragmentation in DHR escalated from 1993 till 2011, the greatest being in the period from 2004 to 2011. One formidable reason was the political instability in the country over the years 1996 to 2007 (Maoist insurgency period) when personnel from the field office along with security check post moved to the city [65–67] seeking security, creating more opportunity for illegal activities in the reserve. However, after the end of Maoist insurgency, negative activities have constrained with appropriate conservation activities [42]. A similar trend has also been observed in Kailash Sacred Landscape of Nepal during the same time period [15].

It is apparent that local people mostly depend on the forest for subsistence needs [68, 69], so sustainable methods should be introduced which can perpetually meet the people’s needs and also protect the reserve at the same time. Establishment of a buffer zone around the reserve just like in other protected areas of Nepal is a plausible way to minimize the dependency of local people on the reserve. Also, upgrading poverty reduction approaches in supporting poorer families in asset accumulation and undertaking alternatives for higher remunerative livelihood strategies will eventually reduce the pressure on environment [70].

## Conclusion

This study accessed spatial and temporal change patterns of land cover and forest fragmentation of DHR between the years 1993 to 2018. The LC change analysis was done for five-time points using a supervised classification method to find out that the forest has increased over the time period throughout the study area while grassland and agriculture & built-up areas have decreased.

The fragmentation process analysis was done with fragmentation spatial model and class metrics was accessed through FRAGSTATS. The multi-temporal overview of forest fragmentation showed that the fragmented patches are concentrated around the forests near human settlement. Shrinkage was the most dominating spatial process for fragmentation followed by subdivision. Most of the expansion occurred around the shrinkage or the subdivision process of preceding years. The high dependency of local people on the forest seems to be the dominant cause of forest fragmentation and forest cover change. The forest fragmentation in different blocks of DHR will have a negative impact on biodiversity so appropriate strategies to control forest shrinkage and forest subdivision are required to be executed. There should be a clear mechanism for resource sharing like creation of a buffer zone area around the reserve and promotion of poverty reduction approaches. This study has contributed to filling up the information gap in a poorly researched Himalayan area with poor data availability. Such information will help develop conservation schemes, plan prioritized reserve management and make policies.

## Acknowledgement

We are thankful to local elders and staff of Dhorpatan Hunting Reserve for providing historical and current information where the researchers couldn’t have direct access.

